# Explanatory latent representation of heterogeneous spatial maps of task-fMRI in large-scale datasets

**DOI:** 10.1101/2021.03.10.434856

**Authors:** Mariam Zabihi, Seyed Mostafa Kia, Thomas Wolfers, Stijn de Boer, Charlotte Fraza, Sourena Soheili-Nezhad, Richard Dinga, Alberto Llera Arenas, Danilo Bzdok, Christian F. Beckmann, Andre Marquand

## Abstract

Finding an interpretable and compact representation of complex neuroimage data can be extremely useful for understanding brain behavioral mapping and hence for explaining the biological underpinnings of mental disorders. Hand-crafted representations, as well as linear transformations, may not accurately reflect the significant variability across individuals. Here, we applied a data-driven approach to learn interpretable and generalizable latent representations that link cognition with underlying brain systems; we applied a three-dimensional autoencoder to two large-scale datasets to find an interpretable latent representation of high dimensional task fMRI image data. This representation also accounts for demographic characteristics, achieved by solving a joint optimization problem that simultaneously reconstructs the data and predicts clinical or demographic variables. We then applied normative modeling to the latent variables to define summary statistics (‘latent indices’) to find a multivariate mapping to non-imaging measures. We trained our model with multi-task fMRI data derived from the Human Connectome Project (HCP) that provides whole-brain coverage across a range of cognitive tasks. Next, in a transfer learning setting, we tested the generalization of our latent space on UK Biobank data as an independent dataset. Our model showed high performance in terms of age and predictions and was capable of capturing complex behavioral characteristics and preserving the individualized variabilities using a highly interpretable latent representation.

## Introduction

One ultimate challenge in the application of machine learning to neuroimaging is to find an optimal summary of the complex spatial information encoded in brain images into biologically interpretable representations which can be used to understand inter-individual differences, learn associations with cognitive variables and to discover biomarkers that explain the biological underpinnings of healthy and disordered mental states^1–5^.

Neuroimaging studies have traditionally had a limited number of high-dimensional datasets, which until recently had hindered employing complex deep neural network models for a time due to the curse of dimensionality^6^ The recent increase in the availability of large-scale neuroimaging data has provided a great opportunity to move toward employing complex nonlinear methods, for example based on deep learning approaches^7–13^. Many deep learning studies in neuroimaging use hand-crafted features ^5,14–17^e.g., regions of interest (ROIs) or image-derived phenotypes (IDPs), which are potentially suboptimal for prediction because (i) hand-crafted features may not accurately capture complex structural or functional brain characteristics e.g. overlapping latent representations encoded in the brain, nor their intricate relationships with behavior and (ii) they do not benefit from the strength of deep neural networks in automatically learning the optimal representation from the data (for example using convolutional filters). Particularly in task fMRI studies, which are designed to study mappings from brain activations to cognition and behavior, there are many challenges in understanding the underlying mechanisms, including the extensive heterogeneity across subjects, finding an optimal representation, and a reliable reference to compare the activations ^18–24^. Consequently, using hand-crafted features potentially leads to losing information relevant, for example, for understanding inter-subject variability ^5,25^. In these scenarios, learning an optimal representation of high-dimensional neuroimaging data rather than – for example – using pre-defined ROIs may enable us to better understand individual variation and more accurately predict clinical and cognitive variables. This representation, also called a latent representation, allows us to reduce the data dimensionality and extract only the essential features from the data. In other words, a latent representation model maps complex and high-dimensional data into a reduced and low-dimensional space^26^.

Having learned the latent representation, we suggest that there are two steps to assess the latent representations: first, whether the derived latent representation shows a stronger association with cognitive, clinical and demographic variables, here referred to collectively as ‘non imaging-derived phenotypes’ (nIDPs) compared to data in the original space (e.g., mapping from raw image data or hand-crafted features to behavioral scores) and further, whether the latent space can be generalized to accurately reconstruct or make predictions for new data (new brain scans, new participants or new scanning sites) which may have a partially different distribution. In the event that this is proven applicable, then, the knowledge learned from one large-scale dataset can be transferred to modeling smaller datasets in a transfer learning paradigm ^27^.

Most applications of deep learning in neuroscience focus on learning a latent representation that is optimized for a single supervised learning problem, such as predicting age or sex (e.g. 11 ^7,28,29^). However, this may reduce the generalizability of the learned latent representation to other problems. Therefore, we sought to learn a general-purpose latent space that is not bound to a particular task, and instead aims to learn features from the data that are predictive of many different cognitive scores. There have been a number of efforts to that end, e.g. to generate synthetic neuroimaging data ^30–33^. However, most of these studies evaluate the data representation on the basis of specific measures like reconstruction error. However, this does not necessarily suggest that the latent space presents relevant features, and what is more important is how accurately such representations can associate with nIDP measures. Although linear data-driven transformations like Principal Component Analysis (PCA) and Independent Component Analysis (ICA) ^34–38^ are widely used for feature representation and dimensionality reduction in neuroimaging, these methods often fail to extract complex nonlinear relationships in data. ^39,40^

In this paper, we propose to explore the value of learning a general purpose nonlinear latent space representation of task-fMRI images using a 3-dimensional semi-supervised autoencoder (AE). Autoencoder neural networks provide a powerful tool in various applications in neuroimaging studies, from image segmentation to abnormality detection and latent representation ^8,9,41–45^. Complementary to existing approaches, we are interested in automatically learning contextual features using an autoencoder. In addition, we show how we can control the latent representation learned by the autoencoder by adding a supervised learning term to the reconstruction (i.e. in a joint optimization framework). Briefly, an autoencoder is a deep neural network architecture that consists of two parts an encoder and a decoder. The encoder projects the inputs to a lower-dimensional latent space using a non-linear transformation. The decoder translates back the latent space to the original space by reconstructing the inputs^46^. Here we controlled the search space by adding age and sex to the loss function minimized by the model. In contrast to many previous approaches, this does not require the prior specification of nodes or regions of interest, can learn overlapping representations, can use the full range of spatial patterns in the fMRI signal and takes advantage of the strengths of deep learning, for example by learning convolutional filters that capture low-level features of the images.

More specifically, in a fully data-driven approach shown in Figure 1, we showed that there is useful information about the data in the nonlinear latent space that is not fully captured by a linear data representation and that such information can be extracted using a hierarchical non-linear autoencoder architecture with joint optimization with age and sex prediction. Here, we employed an autoencoder with an architecture designed from the ground up for task-fMRI data and provide a method for visualizing, exploring and interpreting the learned representation. Last, to illustrate how this model can be used to understand inter-individual differences we applied normative model^47–49^ on the UMAP of latent variables to separate variation in that is principally age-related (encoded by the normative model) from inter-individual differences that manifest as deviations from an expected age-related pattern (encoded in the deviations of the normative model). We these use these deviations for detecting associations with nIDPs. We trained our model with multi-task fMRI data derived from the Human Connectome Project (HCP)^18^ that provides whole-brain coverage across a range of cognitive tasks. Next, in a transfer learning setting, we tested the generalization of our latent space on a UK Biobank dataset^22^ after fine-tuning. Our experimental results show that our nonlinear data representation provides a strong foundation for subsequent analysis of brain-behavior mappings and results in strong associations between our latent index and unseen nIDPs.

**Figure 1:**
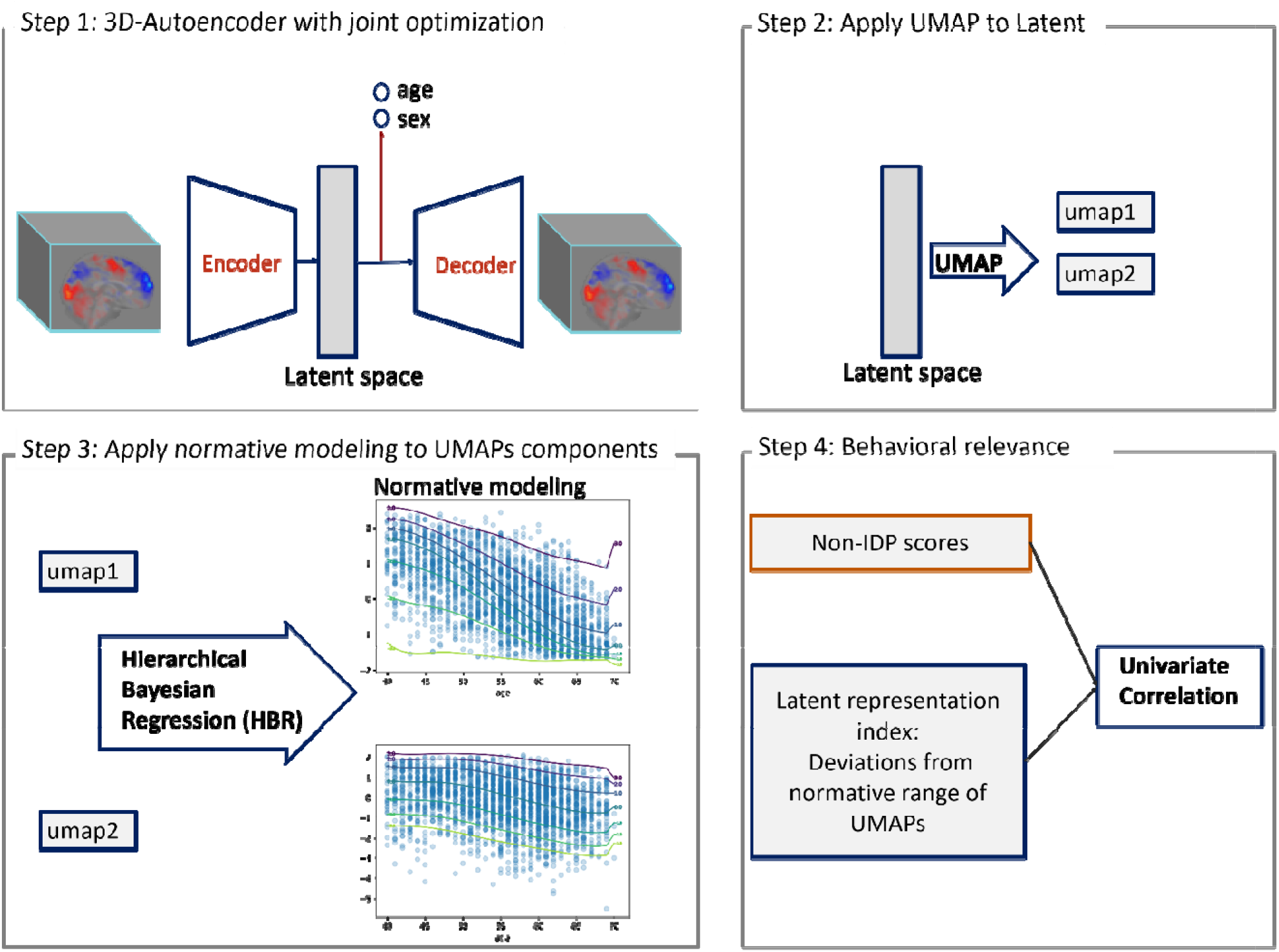
Method overview: step1) training semi-supervised AE model with joint optimization of age and sex prediction. Step2) applying UMAP transformation to the latent variables of semi-supervised AE. step3) applying HBR normative modeling to the components of UMAPs. Step4) measuring the correlation of non-imaging scores (behavioral, cognitive and clinical scores) and the deviation value from normative range of UMAP components (latent representation index)

## Methods

### Data

Two different datasets were used in this study. This first dataset consists of task-based fMRI data from the HCP ^18^ S500 release. The second tfMRI dataset is from the 2020 UK Biobank imaging release^50^.

#### HCP

We used tfMRI contrast data from 468 participants in total (187 males and 281 females, Age= 29.2±3.5) from seven different tasks (emotion processing, gambling, language, relational processing, social cognition, motor, working memory) across 86 contrasts which served as the basis in previous brain-imaging work ^51,52^. This yields a total of N_≈_40K task-fMRI scans. The HCP dataset is well suited for this purpose because the task battery covers a wide range of cognitive domains and the neuronal activations associated with the task provide good coverage of the entire brain ^19^. The number of participants may vary from task to task; not all the participants have data in all the tasks. While HCP has a large number of samples, the number of participants is relatively small. Therefore, we split data into 5 subsets in a 5-fold cross-validation scheme. The splits are made carefully at the subject level so that each fold contains all the contrasts for a specified set of subjects in order to prevent overly optimistic estimates of generalizability due to the correlations between different contrasts from the same subject. More specifically, in each fold, about 95 participants (20% of the data) were reserved for the test set (N=8K brain scans) and the rest for the training (N= 32K brain scans, 373 participants). For each fold, we trained a separate model. Moreover, to further guard against overfitting, an independent set of subjects were used to determine the optimal model architecture (see below and in the supplementary material for details).

#### UK Biobank

We used UK Biobank task-fMRI contrast data from 20781 participants and 5 contrasts, in total N_≈_104K scans (9,860 males, 10,921 females, Age=54.6 ±7.4). The fMRI data derived from UK Biobank uses the same paradigm as the emotion task from the HCP with only minor modifications (e.g. to accommodate shorter run length) ^22,50^. Since UK Biobank provides a larger number of participants than HCP, we trained separate model for each contrast. We randomly selected N=15585 of participants for the train set and 5196 for the test set. All the contrast-models employ the same dataset configuration (the test and train sets).

#### Non-imaging data

The UK Biobank study provides an extensive number of clinical, behavioral, lifestyle and cognitive scores, which we categorized to seven groups e.g., cognitive phenotypes, lifestyle, and mental health (see supplementary information for the full list of categories). We only included the measures that their scores are available more than half of participants. Moreover, in line with previous studies ^53,54^ the measures that had same value for more than 80% of the participants were excluded from further analysis..

### Preprocessing image

For both datasets we used the volumetrically preprocessed images in standard reference space provided by the respective consortia ^55,56^ (for HCP using the ‘minimally processed’ pipeline ^55^). Subsequently the scans images were downsampled from 2mm to 3mm voxel resolution to reduce the computational burden then cropped tightly to the whole brain such that the dimension of the image decreased to 56×64×56. The model was trained on the whole-brain contrast images.

### Model architecture

We developed a deep 3D-convolutional autoencoder that learns to encode and decode task-fMRI images using HCP data. Since there are many choices that need to be made regarding the architecture of the autoencoder, we performed a pilot study on a subset of data that were discarded before fitting the final model. Here, we selected the architecture for the autoencoder using held out data (N=30 participants reserved data, scans_≈_2580). Full details of this procedure are provided in the supplement. The final architecture was as follows: Each encoder and decoder of the semi-supervised AE had three hidden convolutional layers with 3×3×3 kernel size. The bottleneck of the model is a dense layer contains 100 nodes. Each layer except the output layer were follows by RELU activation function^57^ to add non-linearity and sparsity to the network and to reduce the likelihood of vanishing gradient. The output layer was followed by linear activation function. To increase the robustness of the model and avoid overfitting, we added drop-out^58^ (drop-out level=.2) to each layer except the output layer. To avoid the risk of a degenerate solution, where the autoencoder simply learns the identity function, we added Gaussian noise^59^ (mean=0, standard deviation =0.1) to the input layer to randomly corrupt the data (see supplementary for details about the optimization of the architecture of the semi-supervised AE).

The loss function to train the model contains two parts; an unsupervised and a supervised loss. The supervised loss simply is the mean squared error of reconstruction image of noisy image and the original image. The supervised loss incorporated into the control of latent space of the autoencoder; Here, we added age and sex as supervised part of calculating the loss function. We used age as a continuous variable rather than a one-hot encoded matrix (i.e. which would effectively treat the regression as a classification problem^60^). This enables us to generalize beyond the age range used in the training dataset, which is important for transfer learning because of potential differences between cohorts. So the training loss is defined by:

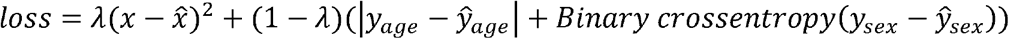

which x is the input image and y_*age*_ and y_*sex*_ are age and sex. The first term refers to unsupervised loss which is the usual autoencoder loss and the second term refers to supervised loss. To balance the supervise and unsupervised loss in terms of scale, we used coefficient *λ* which specifies the importance of supervised loss e.g., *λ*= 1 means completely unsupervised autoencoder (Vanilla AE). We trained our model with different *λs* to select the optimum value in terms of unsupervised and supervised loss.

### Training the model

The training data was normalized with zero mean unit variance across the feature. The layers weights were initialized using Xavier initialization^61^. First, the model was trained using HCP data with 1000 epochs and using Adam^62^ optimizer by adaptive learning rate. The base learning rate was set at .001 and with exponential learning rate decay over each epoch reached 0.0003. Last, the mini_□_batch gradient descent was conducted with the size of 10 images.

Having the model trained by HCP, the network was trained with same hyperparameters again using UKB data as a fine-tunning step. Since the age range is very different across these two datasets, none of the layers were frozen here. Instead, using the same model, the weights of the trained model by HCP used as initial weights and the base learning rate decreased to 0.0003 to train UKB data.

### Latent space representation using UMAP

To visualize and evaluate our model quantitatively, we visualized the latent space using a Uniform Manifold Approximation and Projection (UMAP) approach ^63^ with two components. UMAP is a manifold learning technique similar to t-distributed stochastic neighbor embedding (t-SNE) ^64^ that preserves the local structure of high dimensional data in a nonlinear space. UMAP is superior to tSNE since it better preserves the global structure of data (in addition to its local structure). Furthermore, it is more stable under perturbation or resampling of the data.

Here, to visualize the latent space with two UMAP components, UMAP model was fit using train latent variables without any labeling. To ovoid over-engineering the results, we applied UMAP with the default parameter settings. The size of local neighborhood to learn the manifold structure of the data was set to 15 while the minimum distance of each data in the low dimensional representation was 0.1 in Euclidean distance. Later, this model was applied to the predicted latent variables of test images. We leave further optimization of these parameters for future work.

To assist the interpretation of the latent space, we use a simple method to project back the latent spaces in input (i.e., brain) space. To achieve this, we take advantage of the fact that the UMAP algorithm finds clearly separated clusters for the different fMRI contrasts (see results below). Then, for each contrast, we calculated the center of its cluster (i.e., the centroid of K-means clustering) in 2-dimensional UMAP space. We transformed these centroid points to the latent space (using the inverse UMAP transformation) and used the decoder component of the autoencoder to reconstruct the images corresponding to these cluster centers.

### Associations with nIDPs

#### Normative modeling of UMAP

To assess the biological validity of our latent space, we calculated the linear association between clinical and behavioral measures and the deviations of UMAP reduced latent space for UK Biobank data. However, since the latent variables are related to age and age has a strong association with many cognitive and behavioral scores, we employ normative modeling on the latent space to separate variation in that is principally age-related (encoded by the normative model) from inter-individual differences that manifest as deviations from an expected age-related pattern (encoded in the deviations of the normative model). The normative modeling approach has been used extensively to model heterogeneity in various psychiatric disorders^17,65,66^. Briefly, this approach provides a statistical estimation of the distribution of brain measures along with the deviations from the reference cohort at the level of each individual participant and

We define the ‘latent index’ as a feature that indicates the deviation of normative UMAP of latent variables of each image. First, we applied normative modeling using a flexible generalization of hierarchical Bayesian regression (HBR)^67,68^ to the UMAP of latent variables to remove the linear and non-linear association of age and sex. Importantly, we used a recent generalization of the HBR method that can handle heteroskedastic and non-Gaussian distributions. Age was defined as a regressor and sex as batch effect. (See supplementary for the details of HBR normative model).This way, for each UMP component of each individual, we obtained the deviation or z-score which the so-called latent index. Then, we used the latent index as an indicator of individualized brain activation variability by measuring the associations of the latent index and nIDPs using Spearman measure.

## Results

### Autoencoder performance

As described above and in detail in the supplement, the optimum number of nodes of each layers and the number of layers of semi-supervised AE model was obtained by a pilot study using independent data and resulted 32,16,8 number of nodes for 3 layers of encoder and 8, 16, 32 for decoder, respectively. *λ* was set empirically to 0.05 in order to balance the supervised and unsupervised loss. (See supplementary documents for more details on the architecture of semi-supervised AE and the latent space visualization for different values of lambda). The out-of-sample of model performance is shown in Table 1.

**Table 1:** Model performance

|  | HCP | UKB |
| --- | --- | --- |
| Image reconstruction error (MSE) | 0.26 | 0.16 |
| Age mean absolute error | 3.13 ± 0.09 | 4.84 ± 0.25 |
| Sex prediction accuracy | 81% ± 3% | 89% ± 3% |

### Visualization of latent space

The scatterplot of UMAP components of the autoencoder’s latent variables is shown in Figure 2. For selected contrasts ^19^ in HCP and Face-Shape emotion tasks in UKB. This figure shows how the data points are distributed in the latent space with regard to age and sex. By contrasting the left and right columns of Figure 3A and 3B its clear that: (i) in the vanilla AE (*λ*= 1) age and sex were not reflected in the latent space, and rather the latent space principally reflects differences between different tasks; (ii) in the semi-supervised AE (*λ* = 0.05), age and sex are more clearly evident in the latent space. This is especially evident in UKBiobank, where the age range is wider.

**Figure 2:**
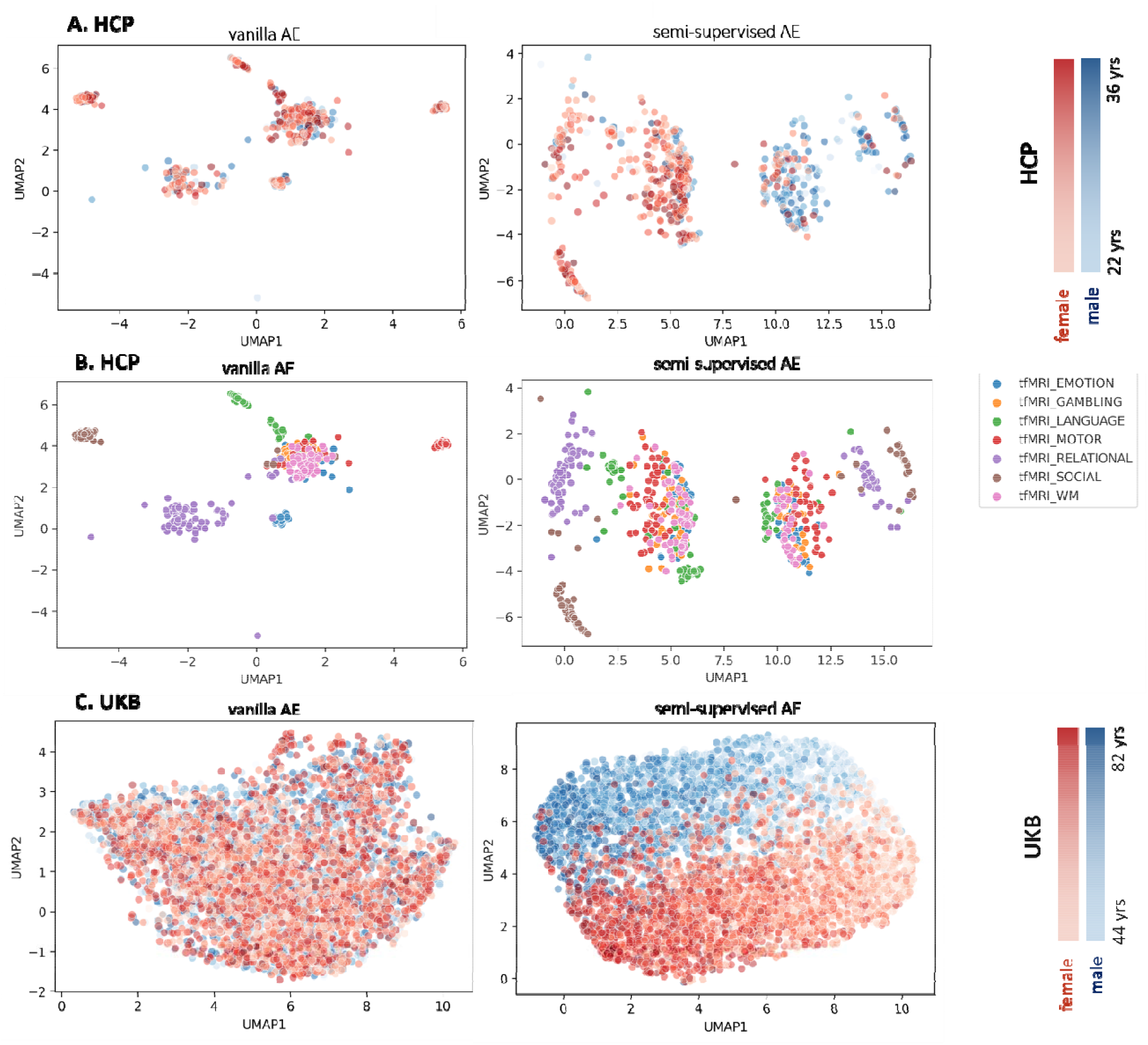
A) UMAPs of latent space of selected contrasts according to Barch 2013 in HCP data in terms of age and sex separation. B) UMAPs of latent space of UCP in terms of task separation. This is identical to panel A, except that the data points are coloured according to task instead of age and sex C) MAPs of latent space of Face-Shape task in UKB data.

**Figure 3:**
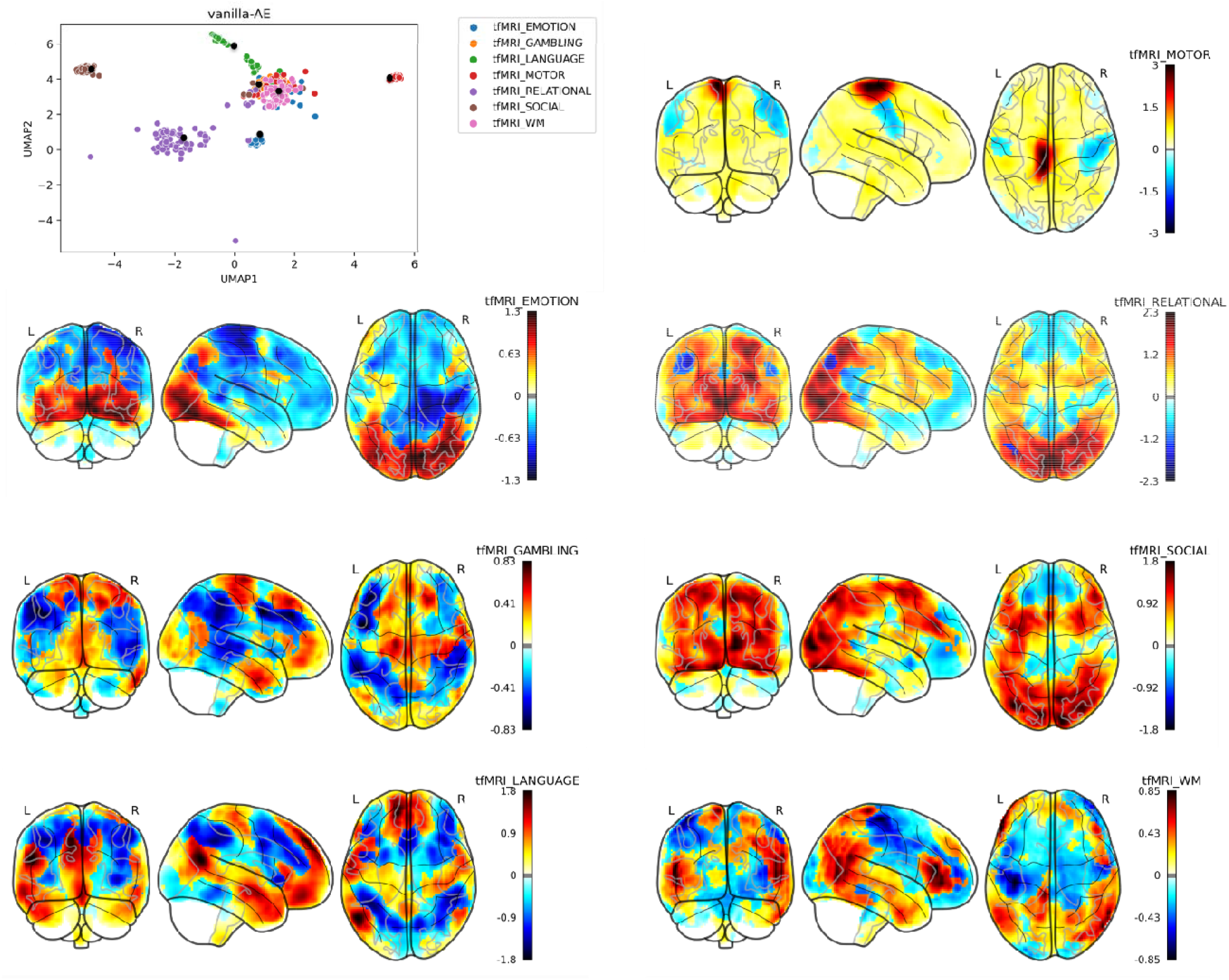
The projection of centroid of UMAP in the latent space to the input brain space. The centers of UMAP of latent space were calculated using K-means clustering across the test data (shown as black points in the panel at the top left). The centroids corresponding to each contrast were passed to encoder of autoencoder to map to input original space.

### Projection the latent representations to brain images

In order to understand relationships in the latent representation in the original space, we show in Figure 3 the centroids of contrasts that are back-projected from the UMAP latent space to the original brain space using vanilla AE. The patterns of activations for these contrasts show an excellent correspondence with the expected task activations as shown in with previous studies (e.g.Barch 2013^19^). For instance, for language task, our projection of latent space to original image space shows the left lateralization which is accord with previous findings in Barch 2013.

### Association between latent variables and non-imaging covariates

The normative models for the UMAP representation of latent variables is shown in Figure 4 (see supplement for measures of fit for the HBR model). In the latent representation the distribution of points has a complex and non-Gaussian distribution, but this can be fit by capitalizing on the flexibility of HBR model.

**Figure 4:**
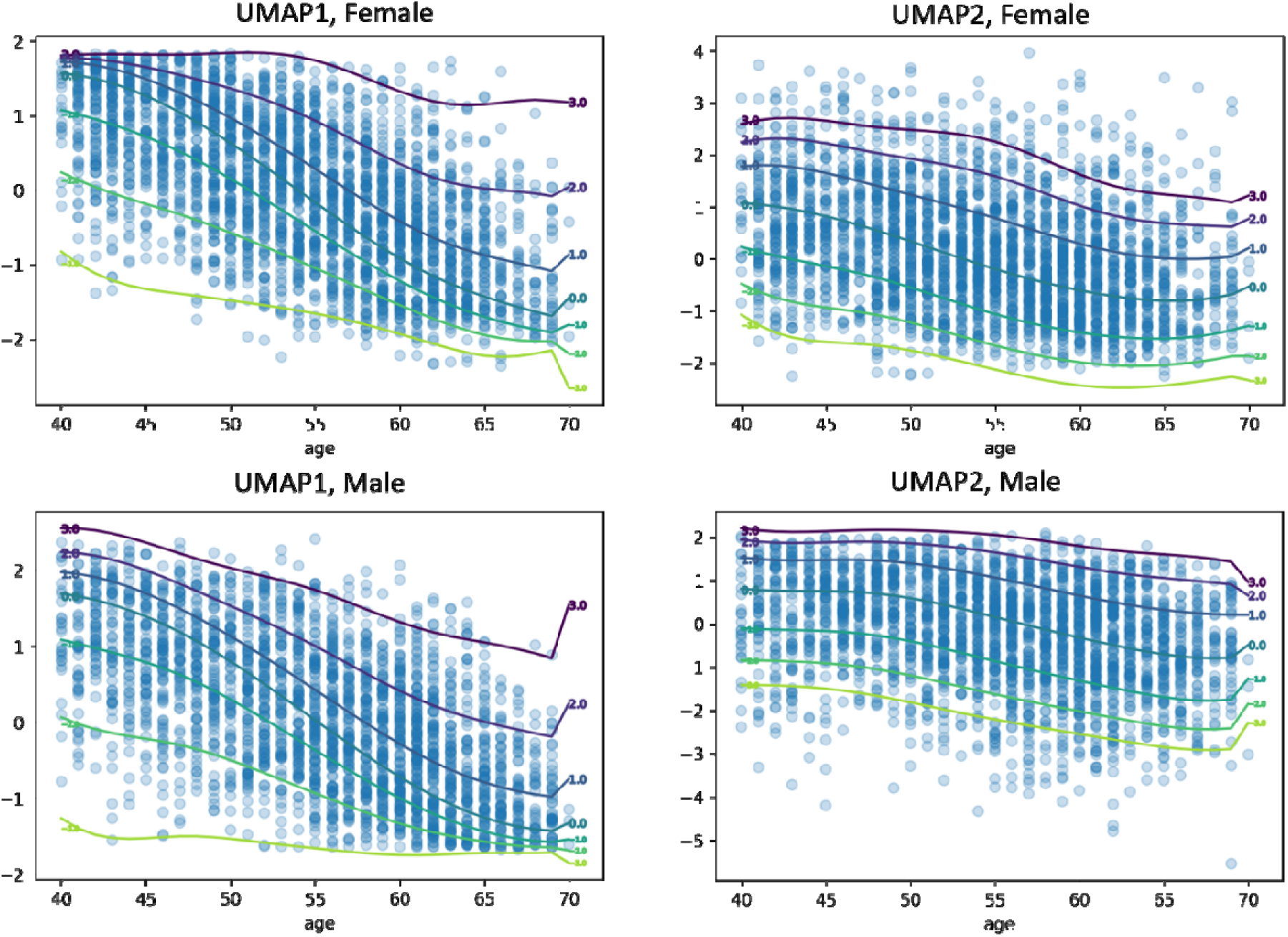
Normative model of the latent space UMAP components. The individualized deviations from the normative range indicates latent representation index.

Figure 5 shows the Manhattan plot of p-value of univariate correlation between non-imaging measures and latent index. This shows that there are strong associations with many nIDPs even after properly accounting for age and sex using the normative model.

**Figure 5:**
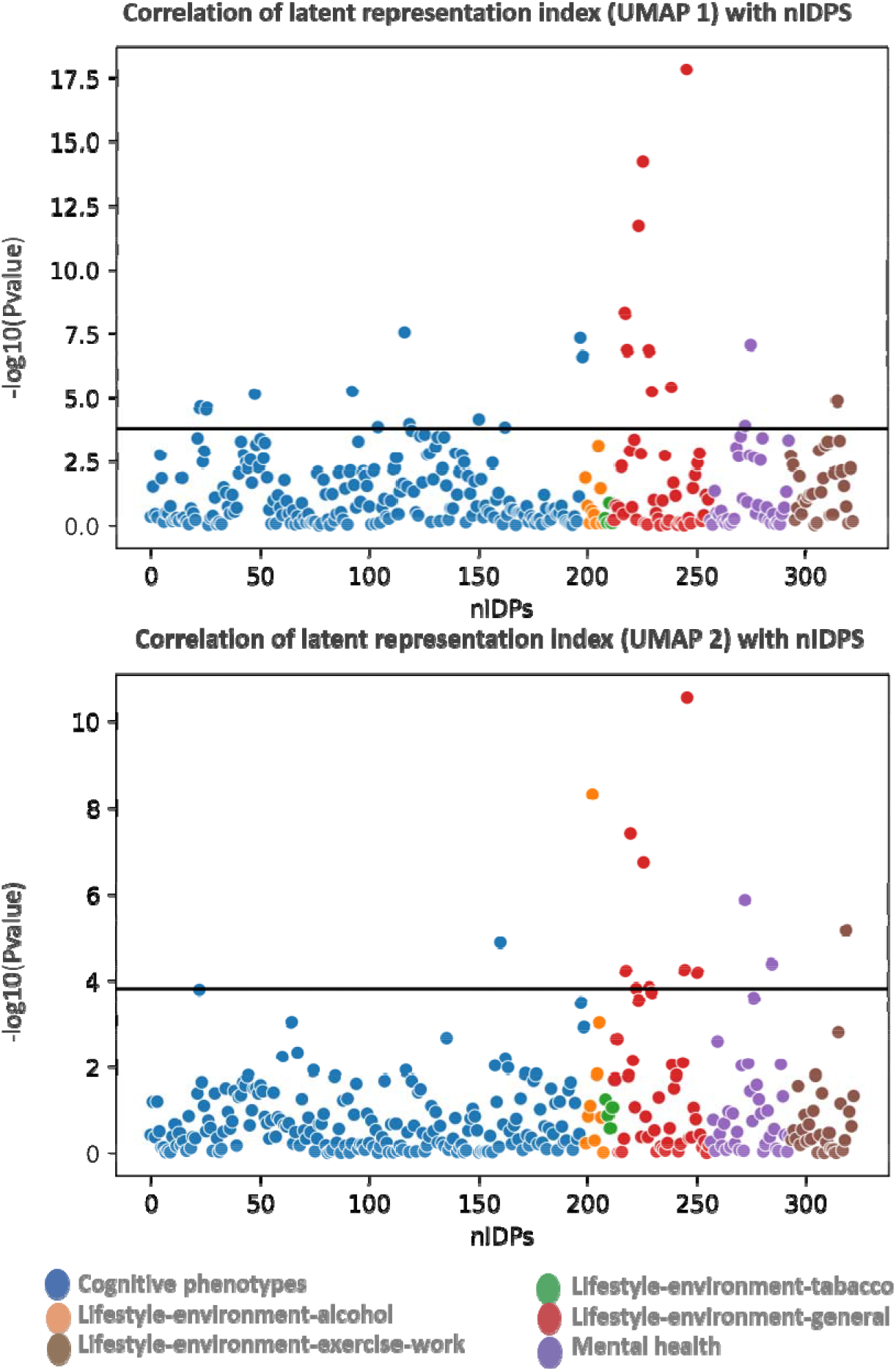
Manhattan plot of p-value of univariate correlation of non-imaging measures with the individualized deviations from normative UMAPs od latent space(latent representation index)s. The black line is Bonferroni-corrected p-value threshold

## Discussion

In this study, we developed a 3-D convolutional autoencoder architecture for non-linear transformation of fMRI data to a low-dimensional, yet informative latent space that allows accurate reconstruction of the data whilst also representing demographic variation. We presented methods to visualize, interpret and control the learned latent space representation and defined a latent index to find a mapping to behavioral measures. We showed that our model learned not only salient features that capture age and other sources of population stratification but are also associated with clinical and behavioral features. Finally, we show that this representation was highly generic and generalized to the UK Biobank population cohort as an independent dataset.

### Learning a generic latent representation

The HCP task-fMRI data enabled us to estimate a generic latent space representation across diverse cognitive tasks^18,19^, whilst also providing good whole brain coverage across all the tasks ^19^. During the training, this mapping allows the autoencoder to learn the various activation patterns across the brain instead of learning specific task-related effects that may be localized to particular brain regions. To validate the generalizability of this latent representation derived from HCP, we used UKB. Complementarily, UKB contains the Hariri faces-shapes emotion task ^69^, which is very similar to the emotion task of HCP (effectively a shorter version). The common contrasts provide a great opportunity for further validation of the model and test the across-cohort generalization of the latent space.

### Mapping the latent space

Since the number of test participants are limited in each model of HCP (N_≈_95) and the age range is limited, the effect of age and sex in the latent space is not clear while the UMAP of UKB generates a clear age continuum and good separation in terms of sex. This indicates that moving from one point in manifold to another can be traced back meaningfully through the input space.

By adding age and sex to the model, we provide a method to explore the functional and anatomical manifolds of brain states by controlling what the autoencoder learns. While unsupervised training of the model yields interpretable representations of different tasks, using semi-supervised autoencoder, our representation was able to be tailored to focus on specific differences. We illustrate this by training an autoencoder that simultaneously reconstructs the data, whilst also predicting age and sex. Importantly, this results in an interpretable latent manifold that clearly reflects individual differences related to the representation of demographic variables in the underlying imaging data.

Projection of latent representation to original space: For the majority of the contrasts and particularly language (story-math), social (theory of mind) and relational (relational-baseline), the projection of the center of K-means of latent space to the original scan image space were in line with findings in ^19^.In the context of interpretability of findings, the meaningful projection of the latent space can be viewed as an example of explainable AI in complex models.

### Association of the latent representation index with non-imaging variables

One important aspect of summarizing the complex spatial maps of tfMRI is to preserve the individualized variability. To complement this, these summaries or representations should contain biological information that can be linked to cognitive, behavioral and clinical characteristics. Due to the fact that the latent space here also represents age and sex, and because age is strongly associated with a variety of cognitive and behavioral scores, the correlation of latent variables and nIDPs may disrupted by the confounding effect of age (see supplementary documents for the correlation of UMAPs and nIPDs). To disentangle clinically relevant variation from variation due to age and sex from the UMAP representation, we applied normative modeling based on hierarchical Bayesian regression. Here, the individualized deviations or latent representation index indicates the distance from the normative latent variables transformed by UMAP. We showed that this index is strongly associated with several nIDP scores after accounting for confounding variables (age and sex). Hence, the notion of normative latent variables may provide the basis for the development of a biomarker that predicts cognitive and behavior characteristics.

### Network architecture

The architectural hyper-parameters of the autoencoder were chosen during the pilot study, solely based on how the models performed in terms of the reconstruction error and no other readouts i.e., non-imaging measures were used for evaluations of the models and the data used for the pilot study were not reused. Some decisions about the network structure have been made before estimating the model. For example, to preserve the morphology of the images and hence better interpretability, we decided to use a 3-D convolutional network ^30–32,41^. In order to control order of latent space, we used dense layer in the bottleneck of the autoencoder ^46^.

We emphasize that we designed our autoencoder with the specific nature of our high-dimensional neuroimaging data in mind and therefore, a number of constraints were imposed on the model beforehand. For example, the networks evaluated were not particularly deep, also to reduce the memory usage and computational complexity, we took advantage of the weight sharing of convolutional layers. Here, we are in search of low-level features that may be translation invariant, but a more important benefit is that the weight sharing enables the networks to be scaled to whole-brain data ^70^. The kernel size was set to be 3 × 3 × 3 to keep the details of the downsampled image scans. Average pooling layers were positioned right after each convolutional layer to ignore the sharp features, reduce the number of parameters and consequently, minimize the chance of overfitting. We relied on the pilot study to select the rest of the model’s parameters, such as the number of filters.

Here, we assigned unsupervised (image reconstruction error) and supervised (age and sex prediction) loss function to our semi-supervised AE while the network’s ultimate goal was finding meaningful latent representations of data that can be mapped to the non-imaging variables and interpreted both in the latent space and in the original voxel space. Our model showed high performance in predicting age and sex. The contribution of supervised and unsupervised loss can be also redefined in order to emphasize the optimization process over supervised or unsupervised loss. This results in a semi-supervised setting that allows the latent space to partially encode particular features of the data ^8^. Another interesting future direction is to train an autoencoder to predict different data (e.g., a follow-up timepoint in longitudinal studies). This would serve to sensitize the latent space to changes relevant to ageing or pathology, which suggests that the latent representation may also be useful to generate features for downstream analyses aiming to predict these features.

The increased number of neuroimaging scans provides a unique opportunity to transcend linear mappings, but it is also necessary to acknowledge some limitations. The traditional image processing techniques often used in deep learning are not completely applicable here. For example, while data augmentation using image mirroring, flipping, skewing, or segmenting is a straightforward approach to increase the number of samples and has been applied before in neuroimaging applications ^11^, we did not consider it to be appropriate here because such augmentation strategies do not faithfully preserve invariances known to occur in the brain, for example the lateralization of brain functions e.g. the association of left lateralization in language processing ^71^. Another limitation is computational complexity. In addition, training an autoencoder on large neuroimaging data is computationally more demanding comparing with similar linear models. In this work we set the trade-off parameter (lambda) governing the contribution of supervised and unsupervised loss components in a relatively informal manner since a quantitative evaluation would have required us to define the relative value of each components (e.g. how much to favour prediction of the supervised targets over reconstruction or vice versa). It is possible that more careful optimization of this parameter may yield improved performance where this information can be specified

## Conclusion

Here, we applied 3-dimensional autoencoder to two large-scale datasets to find an interpretable latent representation of high dimensional task fMRI image data by controlling demographic information. We applied normative modeling to the latent variables to define an index to find a mapping to non-imaging measures.

Our model showed high performance in terms of age and sex predictions and moreover, the generalizability of the representation using an interdependent dataset. Last, our model was capable of capturing complex biological, cognitive, and clinical characteristics and preserve the individualized variabilities using a latent representation index.

