## Supplementary for "Explanatory latent representation of heterogeneous spatial maps of task-fMRI in large-scale datasets"

### Supplementary Materials

##### Methods

**Data:**

The age distributions of HCP^1^ and UK-biobank^2^ tfMRI data across sex are shown in Figure S1.

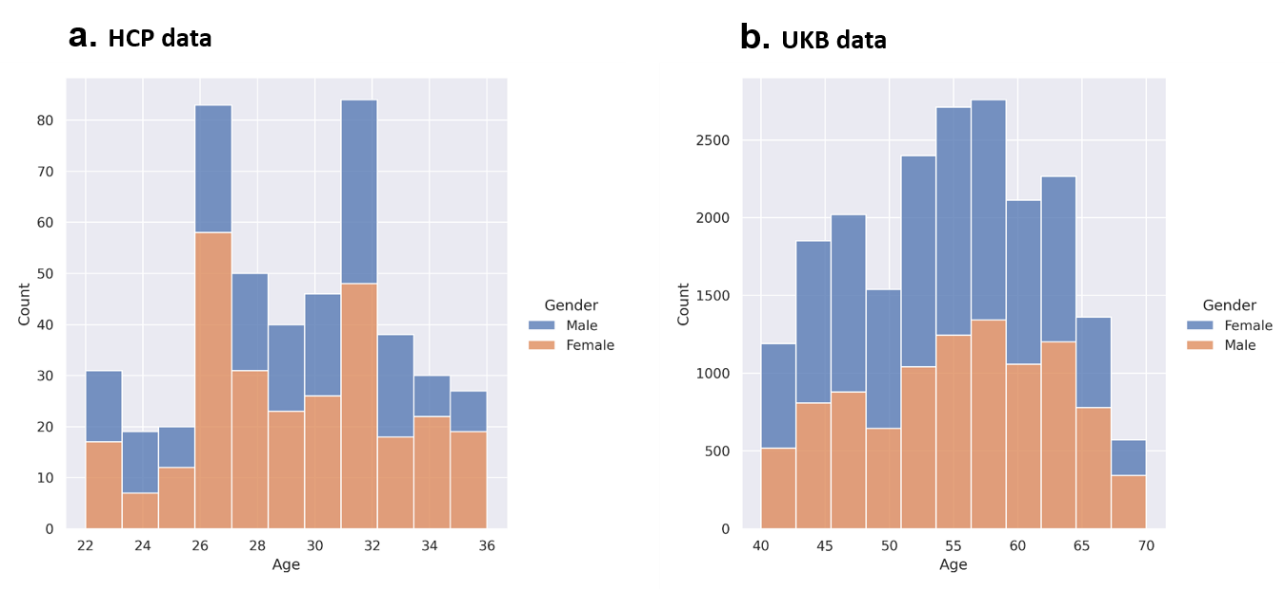

Figure S1: Age distribution in a) HCP and b) UK biobank

The list of UK-bio bank behavioral measures categories ^22^ is shown Table S1.

Table S1: UK biobak non-imaging variable list

|  | **Non-imaging variables categories** |
| --- | --- |
|  | **Basic demographic** |
|  | **Cognitive phenotypes** |
|  | **Lifestyle environmental cohol** |
|  | **Lifestyle environment tobacco** |
|  | **Lifestyle environment general** |
|  | **Lifestyle environment exercise work** |
|  | **Mental health** |
|  | **Age, sex, site** |

Model training:

All the models were trained using NVIDIA P100 GPU, TensorFlow r2.8. The training time for HCP data with N=32K scans was approximately 11.5 minutes per epoch. The fine-tuning time for UKbiobank with N=15K was 2 minutes per epoch. The batch size was 10 for all the models.

#### **Semi-Supervised AE**

The architecture of 3D semi-supervised AE is shown in Figure S2. We used 2580 scans from HCP (N=30 participants) for selecting the architecture and hyperparameters of the network. These scans never used again in the training and test phases.

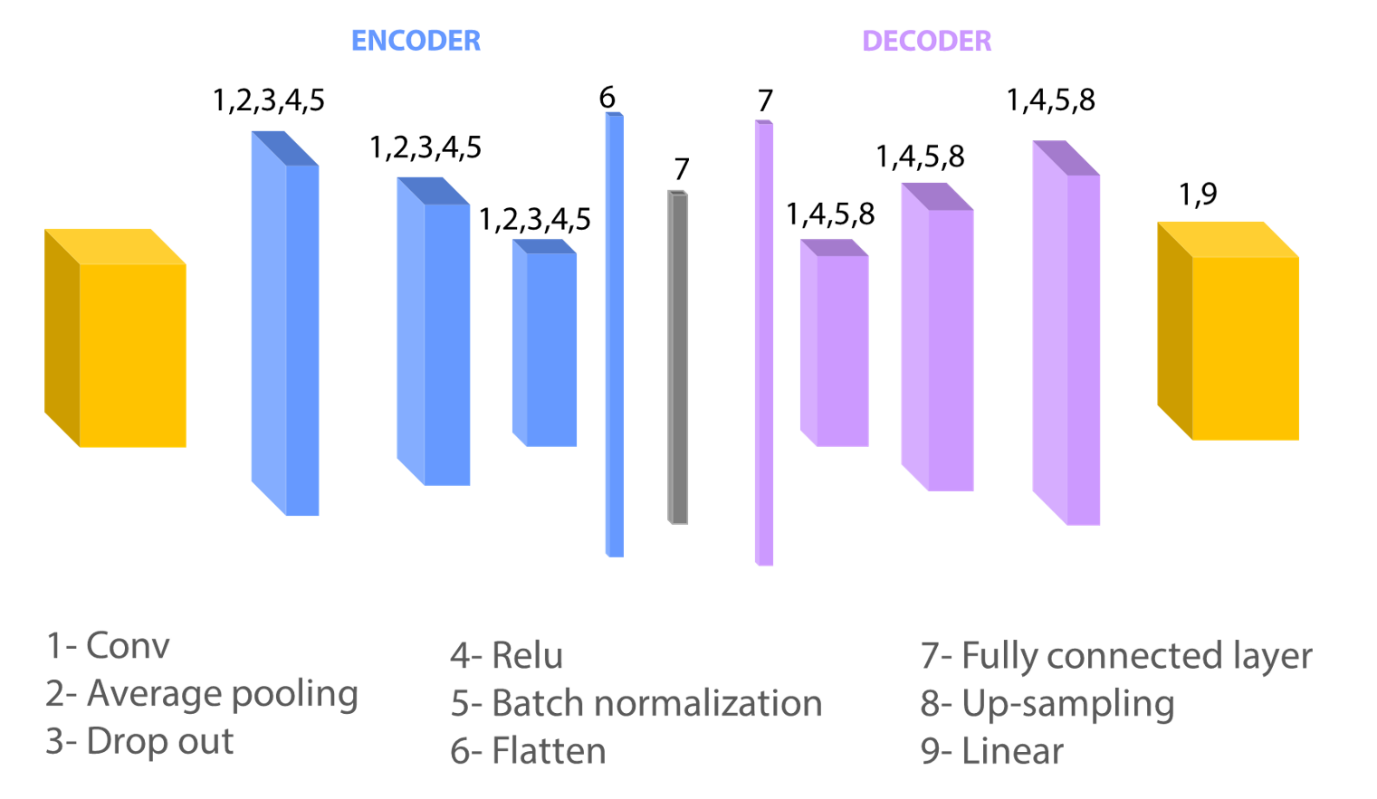

Figure S2: Autoencoder architecture

UMAP hyper-parameters

We tuned the number of neighbors (n_neighbor=15) and minimum distance of the points (min_dist=.1) in a cluster to maintain the balance between the global and local structure of data. Moreover, we selected Euclidean distance for computing the ambient space of data. These parameters provided robust results and better visual separation.

Normative modeling

Briefly, normative modeling presents a probabilistic interpretation of the deviations of reduced latent variables (UMAPs) across all subjects.  To measure the deviations, we used the predicted UMAPs of the normative model for each individual participant, and next converted each to a subject-specific *Z* score as described previously^3^.

We applied Hierarchical Bayesian Regression (HBR) to Normative modeling. Here, we modeled the posterior probability of Y, given X:P(Y|X) using MCMC sampling. For likelihood probability, we used SHASHb distribution, which is a flexible distribution parameterized by corresponding to the mean, variance, skew, and kurtosis respectively. This distribution is very beneficial when the data distribution is non-Gaussian. We used bspline as the basis expansions of the regression. To increase the flexibility of the model, we added random-effect to the mean, variance, skew parameters.

for HBR implementation., we used pcntoolkit (https://github.com/amarquand/PCNtoolkit) framework.

Results

Model selection: Hyper-parameters tuning

Table S2 shows the performance of the models with different configuration and image normalization strategies in terms of normalized root mean square of reconstruction error (NRMSE) which is root mean squared error scaled by the range of the input image.

Table S2: Comparison model performance for different parameters

| nmse | optimizer | dropout | encoder_filters | decoder_filters | lr | scaler_type | dense_layer |
| --- | --- | --- | --- | --- | --- | --- | --- |
| 0.028 | RMSprop | 0.2 | [32, 16, 8] | [8, 16, 32] | 0.001 | Sandard_Feature-wise | 50 |
| 0.033 | RMSprop | 0 | [32, 16, 8] | [8, 16, 32] | 0.001 | MinMax_Feature-wise | 10 |
| 0.027 | **RMSprop** | **0.2** | **[32, 16, 8]** | **[8, 16, 32]** | **0.001** | **Sandard_Feature-wise** | **100** |
| 0.031 | RMSprop | 0 | [32, 16, 8] | [8, 16, 32] | 0.001 | MinMax_Feature-wise | 50 |
| 0.033 | RMSprop | 0 | [32, 16, 8] | [8, 16, 32] | 0.001 | Sandard_Feature-wise | 10 |
| 0.030 | RMSprop | 0 | [32, 16, 8] | [8, 16, 32] | 0.001 | Sandard_sample-wise | 10 |
| 0.032 | RMSprop | 0 | [32, 16, 8] | [8, 16, 32] | 0.001 | Sandard_Feature-wise | 50 |
| 0.028 | RMSprop | 0 | [32, 16, 8] | [8, 16, 32] | 0.001 | Sandard_Feature-wise | 100 |
| 0.030 | RMSprop | 0.2 | [32, 16, 8] | [8, 16, 32] | 0.001 | Sandard_Feature-wise | 10 |
| 0.031 | RMSprop | 0 | [32, 16, 8] | [8, 16, 32] | 0.001 | MinMax_Feature-wise | 100 |
| 0.035 | RMSprop | 0 | [32, 16, 8] | [8, 16, 32] | 0.001 | Sandard_sample-wise | 50 |
| 0.032 | RMSprop | 0.2 | [32, 16, 8] | [8, 16, 32] | 0.001 | MinMax_Feature-wise | 10 |
| 0.036 | RMSprop | 0 | [32, 16, 8] | [8, 16, 32] | 0.001 | Sandard_sample-wise | 100 |
| 0.031 | RMSprop | 0.2 | [32, 16, 8] | [8, 16, 32] | 0.001 | MinMax_Feature-wise | 50 |
| 0.034 | RMSprop | 0.2 | [32, 16, 8] | [8, 16, 32] | 0.001 | Sandard_sample-wise | 10 |
| 0.030 | RMSprop | 0.2 | [32, 16, 8] | [8, 16, 32] | 0.001 | MinMax_Feature-wise | 100 |
| 0.039 | RMSprop | 0.2 | [32, 16, 8] | [8, 16, 32] | 0.001 | Sandard_sample-wise | 50 |
| 0.042 | RMSprop | 0.2 | [32, 16, 8] | [8, 16, 32] | 0.001 | Sandard_sample-wise | 100 |
| 0.032 | RMSprop | 0 | [8, 16, 32] | [32, 16, 8] | 0.001 | Sandard_sample-wise | 10 |
| 0.037 | RMSprop | 0 | [8, 16, 32] | [32, 16, 8] | 0.001 | Sandard_sample-wise | 50 |
| 0.038 | RMSprop | 0 | [8, 16, 32] | [32, 16, 8] | 0.001 | Sandard_sample-wise | 100 |
| 0.033 | RMSprop | 0 | [8, 16, 32] | [32, 16, 8] | 0.001 | MinMax_Feature-wise | 10 |
| 0.037 | RMSprop | 0.2 | [8, 16, 32] | [32, 16, 8] | 0.001 | Sandard_sample-wise | 10 |
| 0.032 | RMSprop | 0 | [8, 16, 32] | [32, 16, 8] | 0.001 | MinMax_Feature-wise | 50 |
| 0.036 | RMSprop | 0.2 | [8, 16, 32] | [32, 16, 8] | 0.001 | Sandard_sample-wise | 50 |
| 0.031 | RMSprop | 0 | [8, 16, 32] | [32, 16, 8] | 0.001 | MinMax_Feature-wise | 100 |
| 0.036 | RMSprop | 0.2 | [8, 16, 32] | [32, 16, 8] | 0.001 | Sandard_sample-wise | 100 |
| 0.032 | RMSprop | 0.2 | [8, 16, 32] | [32, 16, 8] | 0.001 | MinMax_Feature-wise | 10 |
| 0.140 | RMSprop | 0 | [32, 16, 8] | [8, 16, 32] | 0.001 | MinMax_sample-wise | 10 |
| 0.031 | RMSprop | 0.2 | [8, 16, 32] | [32, 16, 8] | 0.001 | MinMax_Feature-wise | 50 |
| 0.162 | RMSprop | 0 | [32, 16, 8] | [8, 16, 32] | 0.001 | MinMax_sample-wise | 50 |
| 0.065 | RMSprop | 0 | [32, 16, 8] | [8, 16, 32] | 0.001 | MinMax_sample-wise | 100 |
| 0.104 | RMSprop | 0.2 | [32, 16, 8] | [8, 16, 32] | 0.001 | MinMax_sample-wise | 10 |
| 0.031 | RMSprop | 0.2 | [8, 16, 32] | [32, 16, 8] | 0.001 | MinMax_Feature-wise | 100 |
| 0.188 | RMSprop | 0.2 | [32, 16, 8] | [8, 16, 32] | 0.001 | MinMax_sample-wise | 50 |
| 0.165 | RMSprop | 0.2 | [32, 16, 8] | [8, 16, 32] | 0.001 | MinMax_sample-wise | 100 |
| 0.084 | RMSprop | 0 | [8, 16, 32] | [32, 16, 8] | 0.001 | MinMax_sample-wise | 10 |
| 0.063 | RMSprop | 0 | [8, 16, 32] | [32, 16, 8] | 0.001 | MinMax_sample-wise | 50 |
| 0.029 | RMSprop | 0 | [8, 16, 32] | [32, 16, 8] | 0.001 | Sandard_Feature-wise | 10 |
| 0.083 | RMSprop | 0 | [8, 16, 32] | [32, 16, 8] | 0.001 | MinMax_sample-wise | 100 |
| 0.116 | RMSprop | 0.2 | [8, 16, 32] | [32, 16, 8] | 0.001 | MinMax_sample-wise | 10 |
| 0.070 | RMSprop | 0.2 | [8, 16, 32] | [32, 16, 8] | 0.001 | MinMax_sample-wise | 50 |
| 0.080 | RMSprop | 0.2 | [8, 16, 32] | [32, 16, 8] | 0.001 | MinMax_sample-wise | 100 |
| 0.028 | RMSprop | 0 | [8, 16, 32] | [32, 16, 8] | 0.001 | Sandard_Feature-wise | 50 |
| 0.028 | RMSprop | 0 | [8, 16, 32] | [32, 16, 8] | 0.001 | Sandard_Feature-wise | 100 |
| 0.030 | RMSprop | 0.2 | [8, 16, 32] | [32, 16, 8] | 0.001 | Sandard_Feature-wise | 10 |
| 0.028 | RMSprop | 0.2 | [8, 16, 32] | [32, 16, 8] | 0.001 | Sandard_Feature-wise | 50 |
| 0.027 | RMSprop | 0.2 | [8, 16, 32] | [32, 16, 8] | 0.001 | Sandard_Feature-wise | 100 |
| 0.029 | RMSprop | 0 | [32, 16, 8] | [8, 16, 32] | 0.01 | Sandard_Feature-wise | 100 |
| 0.032 | RMSprop | 0 | [32, 16, 8] | [8, 16, 32] | 0.005 | Sandard_Feature-wise | 100 |
| 0.034 | RMSprop | 0 | [32, 16, 8] | [8, 16, 32] | 0.0001 | Sandard_Feature-wise | 100 |
| 0.029 | Adam | 0 | [32, 16, 8] | [8, 16, 32] | 0.01 | Sandard_Feature-wise | 100 |
| 0.028 | Adam | 0 | [32, 16, 8] | [8, 16, 32] | 0.001 | Sandard_Feature-wise | 100 |
| 0.052 | Adam | 0 | [32, 16, 8] | [8, 16, 32] | 0.0001 | Sandard_Feature-wise | 100 |

The impact of supervised loss coefficient $\boldsymbol{\lambda}$ on latent

To selection, we trained the model with different values of $\boldsymbol{\lambda}$. Table S3 indicates the impact of $\boldsymbol{\lambda}$ on unsupervised and supervised loss in the small held-out test data set of HCP. As expected, $\boldsymbol{\lambda}=1$ fails to predict age and sex. By increasing the supervised loss impact and decreasing $\boldsymbol{\lambda}$, however, the reconstruction loss increases.

$\boldsymbol{\lambda}=0.05$ shows the better performance in terms of sex and age prediction and reconstruction error.

Table S3: model performance in terms of loss values for different values of λ

| $\boldsymbol{\lambda}$ | Reconstruction error | Balanced accuracy | Mean Absolute Error [years] | Total loss |
| --- | --- | --- | --- | --- |
| 1 | 0,120 | 0,551 | 29,075 | 29,884 |
| 0,995 | 0,125 | 0,966 | 1,275 | 1,564 |
| 0,95 | 0,128 | 0,996 | 0,963 | 1,101 |
| 0,5 | 0,188 | 0,996 | 0,995 | 1,190 |
| 0,05 | 0,193 | 1,000 | 0,820 | **1,016** |
| 0,005 | 0,191 | 1,000 | 1,013 | 1,207 |
| 0,0005 | 0,192 | 0,994 | 0,833 | 1,043 |

Figure S3 Shows how the change of $\lambda$ reflects on latent space in UKB.

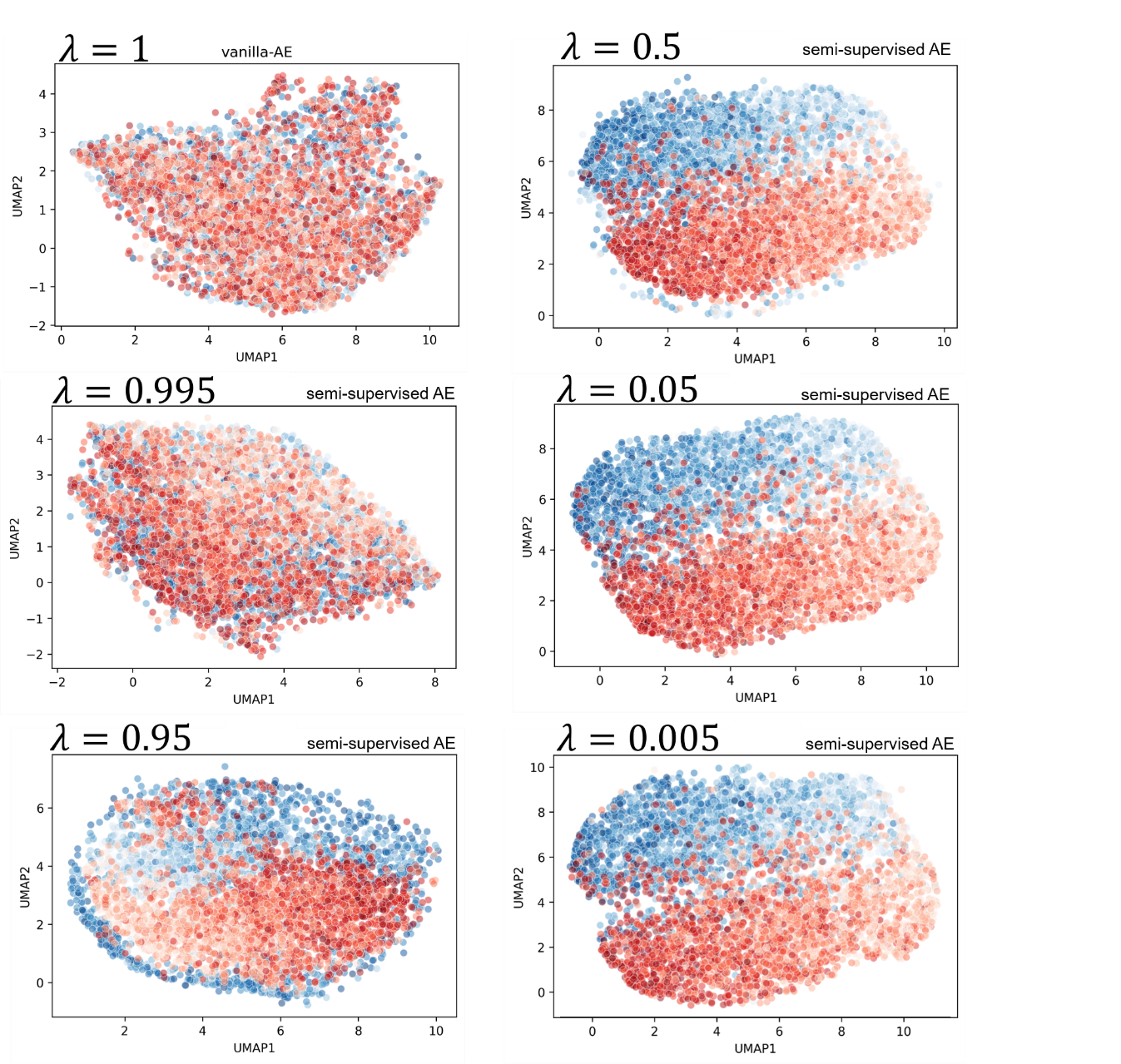

Figure S3: change of latent space representation by changing λ

Autoencoder learning curve

The learning curve of semi-supervised AE is shown in Figure S4.

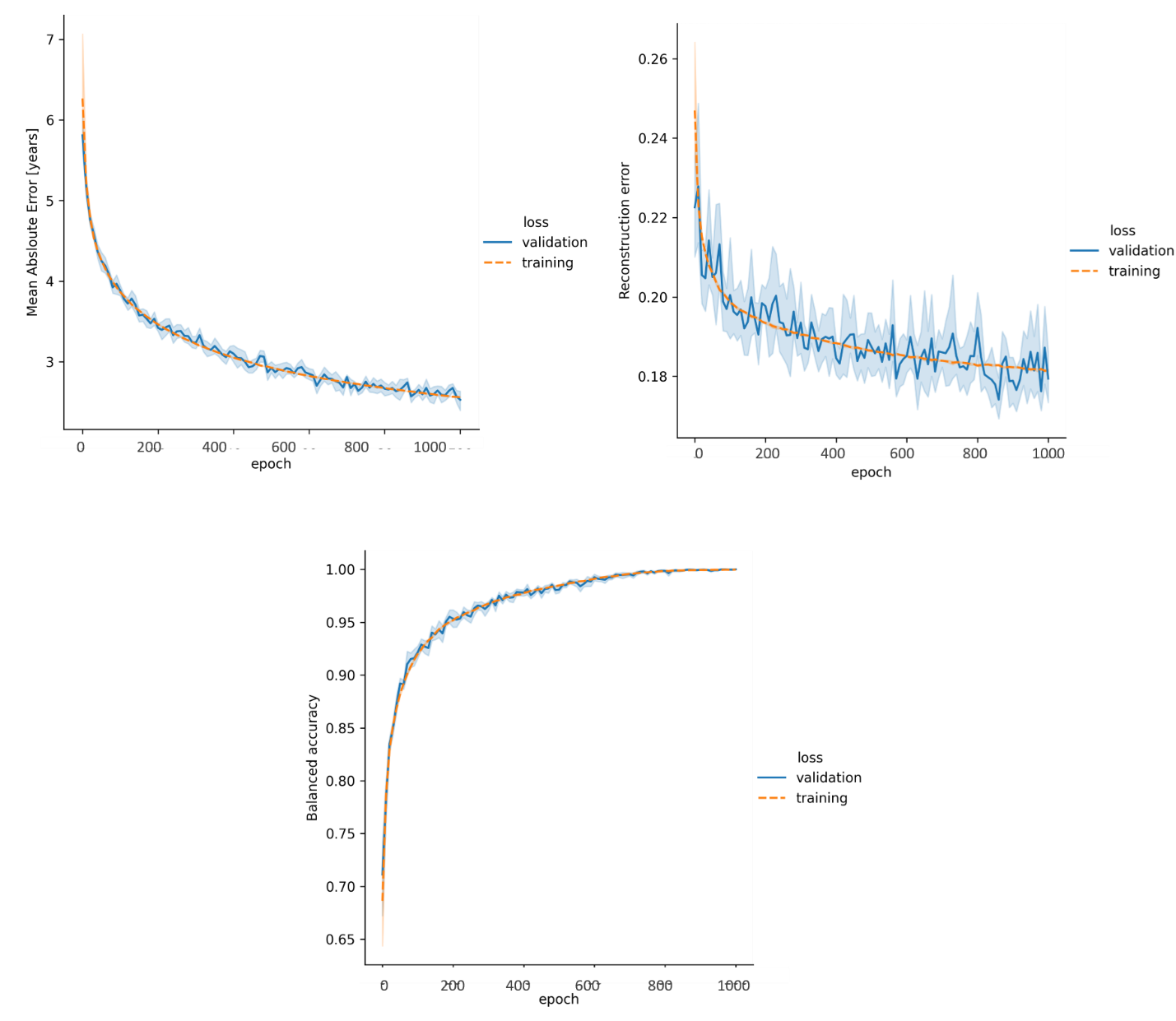

Figure S4: The mean learning curve of semi-supervised AE using UKB

#### **Autoencoder learns different level features in each layer**

Figure S5 shows the weight of kernel in several layers.

Figure S5: Kernels's weights for selected layers

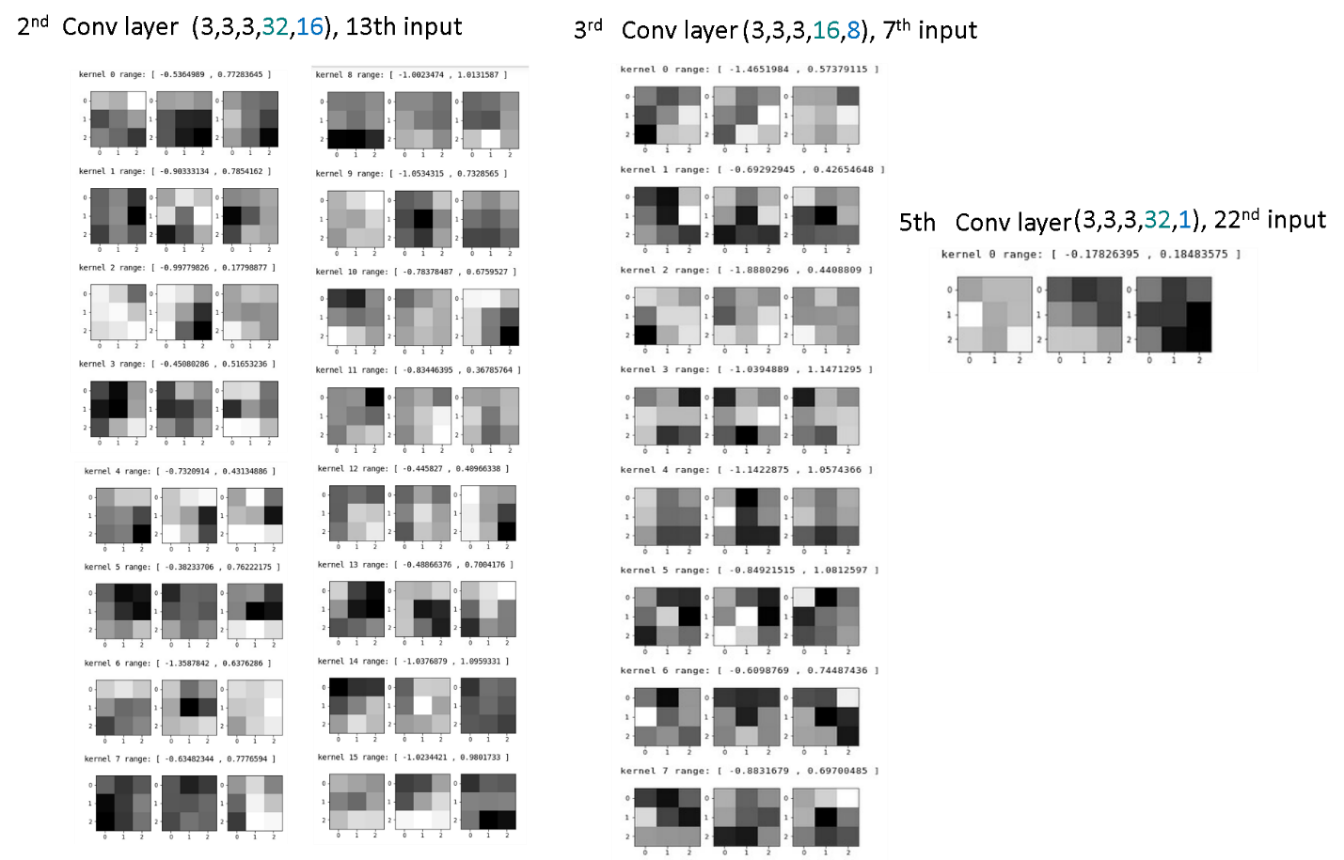

**HBR normative modelling performance**

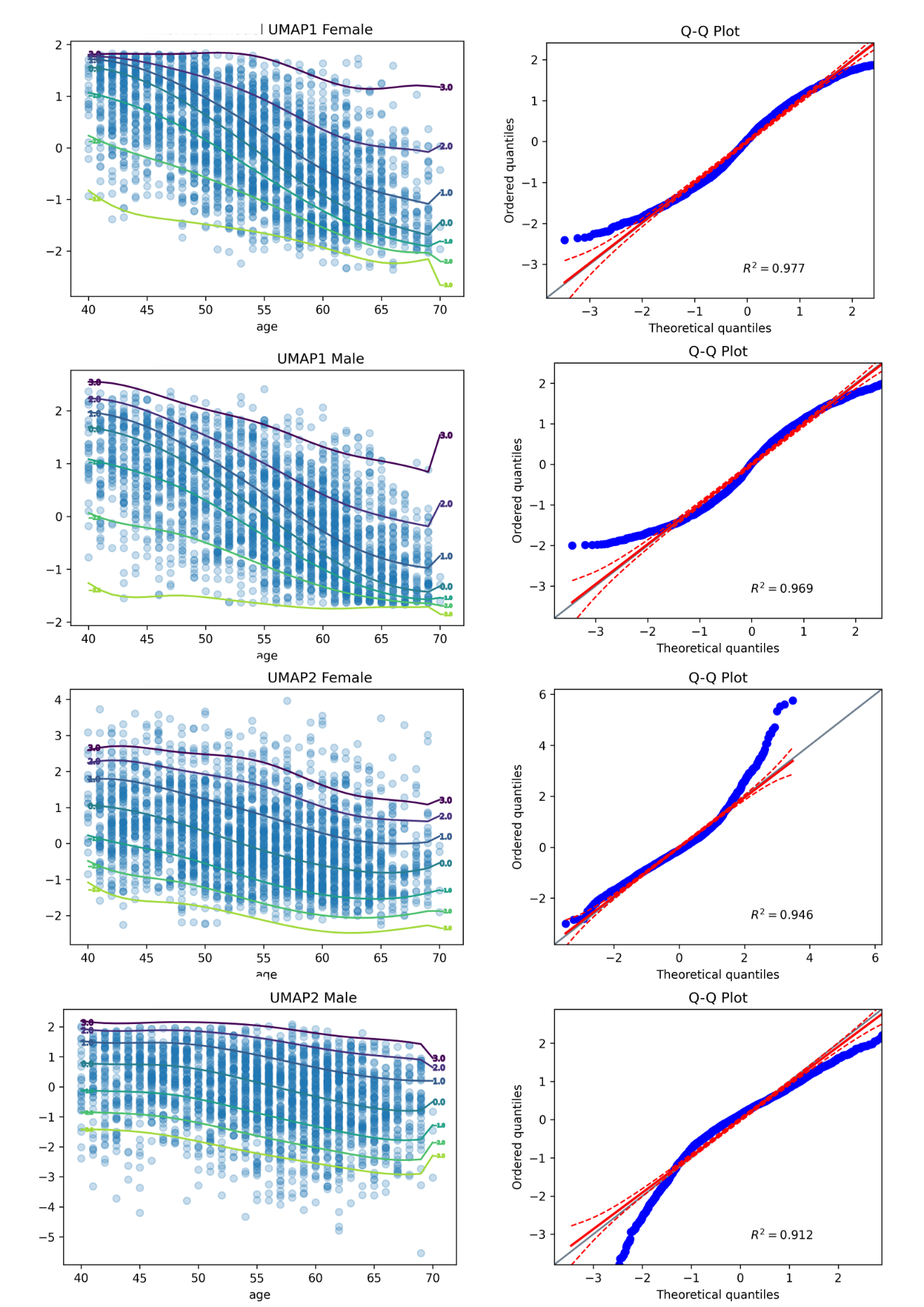

Figure S6: normative UMPAs and qqplot of z-value of normative model

Association of the latent representation with non-imaging variables

This figure indicates that there is highly level of association with age/sex information and other cognitive and behavioral measures. Hence, it shows the necessity of removing the confounding effect of age/sex from the latent variables.

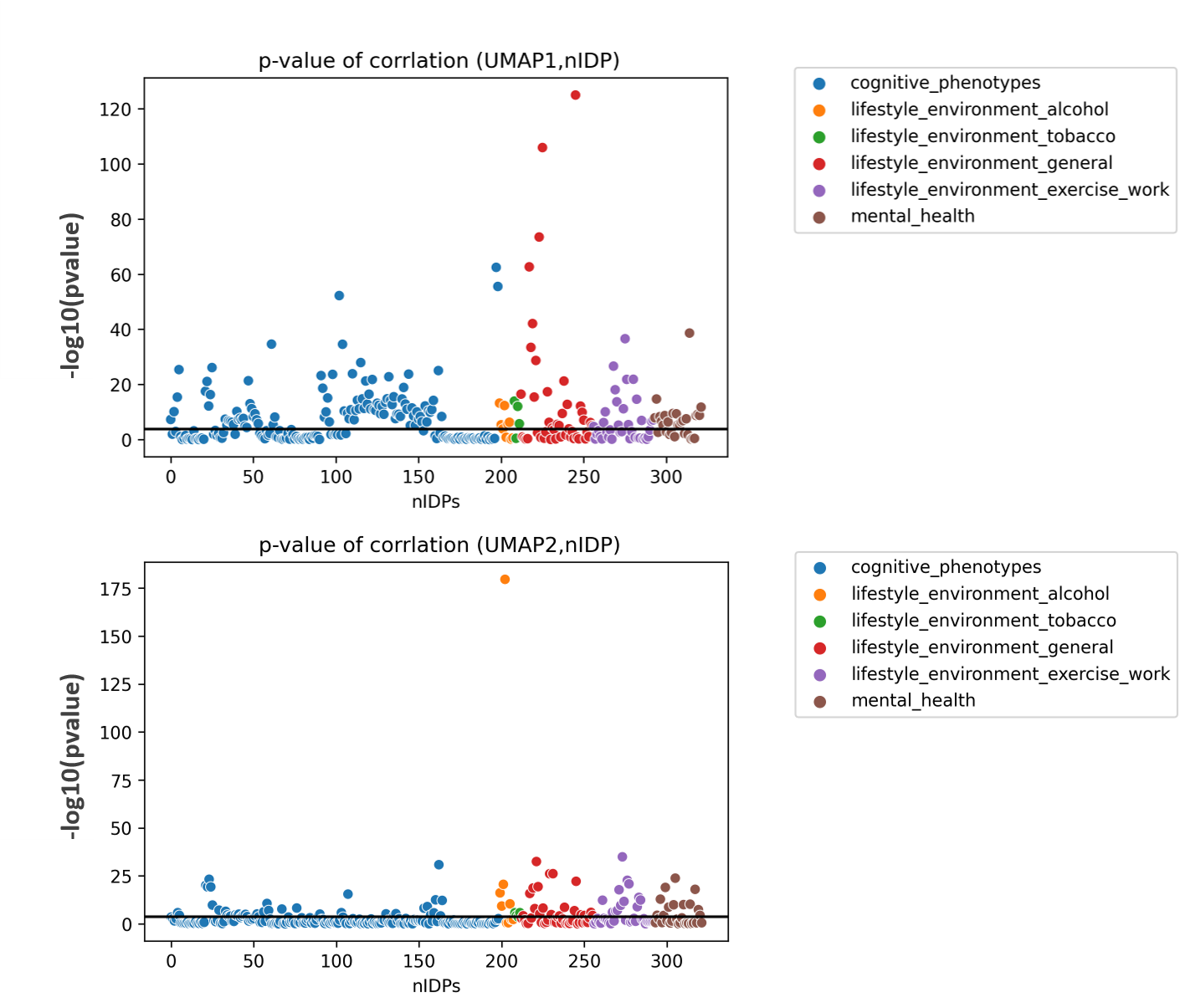

Figure S7: correlation of UMAPs of latent variables with nIDPs
